# Intra-Abdominal Photodynamic Therapy in a Rabbit Model of Perforated Appendicitis

**DOI:** 10.1101/2025.09.24.678338

**Authors:** Timothy M. Baran, Korry T. Wirth, Matthew M. Byrne, Benjamin Gersten, Eric Y. Wang, Darrian S. Hawryluk, Martin S. Pavelka, Nicole A. Wilson

## Abstract

**Importance:** Perforated appendicitis is a common and morbid pediatric surgical condition frequently requiring prolonged antibiotic therapy. Adjunctive intraoperative strategies for local antimicrobial therapy are limited, and photodynamic therapy (PDT) may offer a targeted, resistance-independent approach.

**Objective:** To evaluate the feasibility, safety, and preliminary efficacy of laparoscopic methylene blue photodynamic therapy (MB-PDT) for intra-abdominal disinfection in a rabbit model of perforated appendicitis.

**Design:** Experimental preclinical animal study conducted between 2023 and 2025.

**Setting:** Single-center laboratory-based animal research facility affiliated with an academic medical center.

**Participants:** Nineteen New Zealand White rabbits underwent surgically induced perforated appendicitis with peritonitis via appendiceal ligation and electrocautery perforation.

**Interventions:** Rabbits received laparoscopic MB-PDT (n=9) or control conditions (methylene blue only, no light; n=10) 24 – 40 hours after appendiceal perforation. MB-PDT involved peritoneal lavage with 300 µg/mL methylene blue, followed by 665 nm laser illumination to 25 J/cm^2^. Peritoneal aspirates were collected before and 24h after intervention. Bacterial burden was quantified, and isolated bacteria were submitted to *in vitro* PDT. Tissue samples from intraperitoneal organs were collected 24 hours post-intervention.

**Main Outcomes and Measures:** Primary outcomes included feasibility of PDT delivery, safety (histologic evidence of off-target injury), and preliminary *in vivo* efficacy (change in bacterial burden pre-vs. post-intervention). Secondary outcomes included clinical parameter changes and *in vitro* MB-PDT efficacy.

**Results:** Of the 14 rabbits that developed peritonitis, 10 (age 5–6 months, weight 3.1–3.8 kg) completed follow-up. Laparoscopic MB-PDT delivery was feasible and caused no off-target histologic injury. Bacterial burden increased in MB-PDT animals relative to controls, though not significantly (p=0.18). Post-intervention, body temperature showed a trend toward reduction in MB-PDT animals (−0.37 ± 1.32°C) vs controls (0.26 ± 0.76°C; p=0.27). *In vitro* MB-PDT significantly reduced bacterial burden for all tested species (p<0.001), with greater efficacy in Gram-positive bacteria.

**Conclusions and Relevance:** Laparoscopic MB-PDT was feasible and safe in a rabbit model of perforated appendicitis, with high *in vitro* antimicrobial efficacy. Although *in vivo* bacterial reduction was not demonstrated, these results support further investigation of MB-PDT as a novel intraoperative adjunct for treating intra-abdominal infection.

## INTRODUCTION

Appendicitis is the most common general surgical condition worldwide, and accounts for significant morbidity and health care resource utilization^1,2^. Approximately ⅓ of pediatric cases in the United States present with perforated appendicitis (PA)^3,4^, leading to bacterial spillage into the peritoneal cavity, peritonitis, and often abscess or phlegmon formation. Compared with non-perforated appendicitis, PA is associated with longer hospital stays, higher costs, increased readmissions, and worse postoperative outcomes^3,5–8^.

Antibiotics remain the mainstay of treatment for intra-abdominal infection due to PA, but the growing burden of antimicrobial resistance underscores the urgent need for alternative adjunctive therapies. Photodynamic therapy (PDT), which utilizes light-activated photosensitizers to generate bactericidal reactive oxygen species, is broadly effective against multiple classes of bacteria, including those with antibiotic resistance, and does not have any known mechanisms that lead to acquired antibiotic resistance^9–11^. Methylene blue (MB) has been used successfully as a photosensitizer for antimicrobial PDT treatment of superficial skin and mucosal infections^12–15^, preoperative nasal cavity disinfection^16^, and intra-abdominal abscess treatment^17^.

Although MB-PDT has shown strong *in vitro* efficacy against pathogens commonly found in abdominal infections^18–26^, *in vivo* testing in a representative animal model is a key step toward clinical translation. Rabbits, unlike rodents, possess a robust appendix^27^ and are well suited for modeling perforated appendicitis^28–31^. However, prior rabbit models of PA have struggled to produce a consistently perforated appendix at a predictable time point^28,29^.

In this study, we aimed to: (1) develop a novel, reliable rabbit model of PA with peritonitis, (2) assess the feasibility and safety of laparoscopic MB-PDT in this model; and (3) evaluate *in vitro* MB-PDT efficacy against bacteria isolated from infected animals. We hypothesized that: (1) we could create a consistent rabbit model of PA with peritonitis, (2) MB-PDT would be feasible and safe for intra-abdominal use, and (3) that MB-PDT performed *in vitro* on bacteria obtained from the rabbit model would effectively decrease the bacterial burden associated with PA. A secondary hypothesis was that *in vivo*, laparoscopic MB-PDT would successfully reduce bacterial burden relative to control animals.

## METHODS

After IACUC approval, 19 New Zealand White rabbits (Charles River Laboratories, Wilmington, MA) were purchased at 5-6 months of age (Charles River Laboratories, Wilmington, MA). All rabbits received identical housing, enrichment, and diet. Pain was managed with a multimodal analgesic approach: pre-operative intravenous buprenorphine (0.12 mg/kg), local anesthetic at the incision (1 mg/kg bupivacaine), and transdermal fentanyl patches (12-25 µg/hour). All animals underwent three surgical procedures over three days (Figure 1): (1) induction of PA on Day 1, (2) PDT or control condition on Day 2 with pre-intervention sample collection, and (3) post-intervention sample collection and tissue harvesting on Day 3.

**Figure 1:**
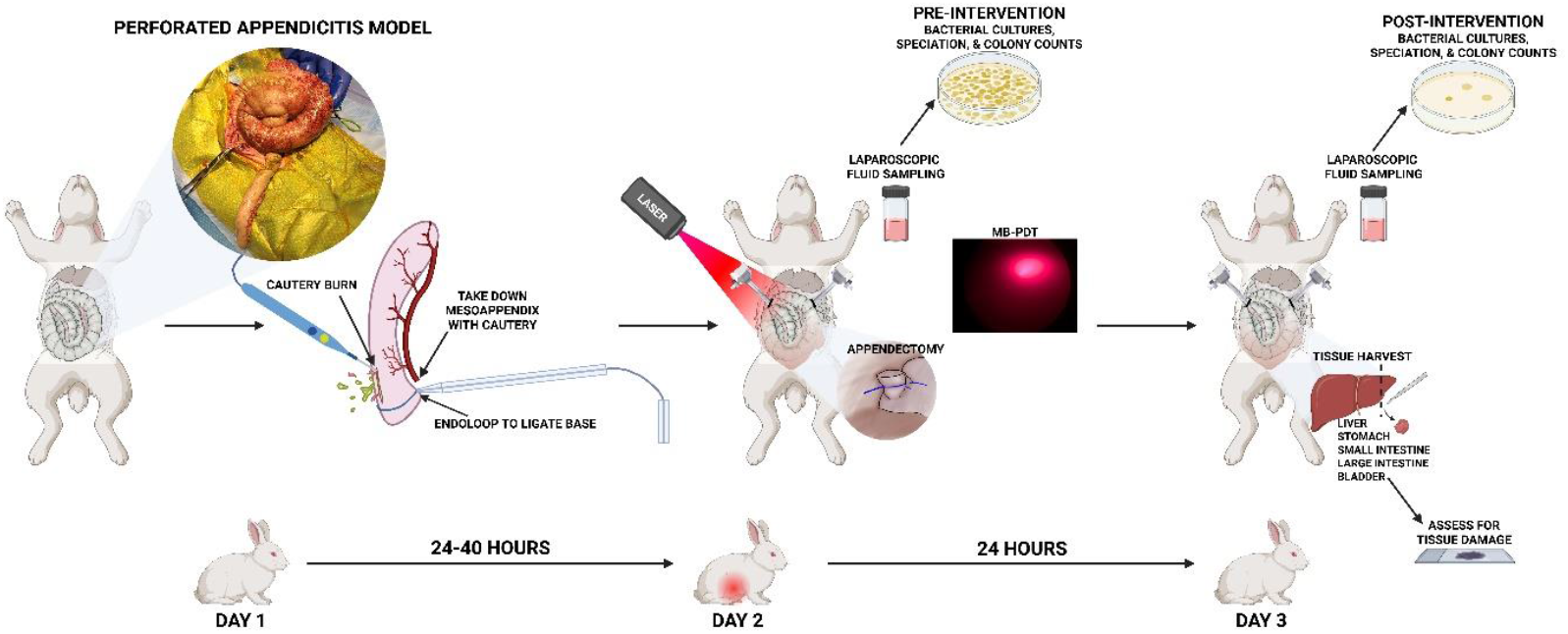
Experimental design. Illustrating induction of perforated appendicitis on Day 1, PDT or control conditions on Day 2 with pre-intervention fluid collection, and post-intervention fluid and tissue collection on Day 3. (Created in BioRender. Wilson, N. (2025), https://BioRender.com/7vr9yjp)

### Induction of perforated appendicitis

The full details of the rabbit model of PA can be found in the Supplemental Material. Briefly, on Day 1, via laparotomy, PA was created via dividing meso-appendix perforator vessels, suture ligating the appendiceal artery, suture ligating the base of the appendix, and creating a full-thickness linear perforation on the anti-mesenteric appendiceal surface using electrocautery (Figure 2b). Of note, appendiceal artery ligation was intentionally not performed in the last treated animal. The rabbit was then returned to the vivarium for 24 hours, during which time PA with diffuse peritonitis developed (Figure 2c).

**Figure 2.**
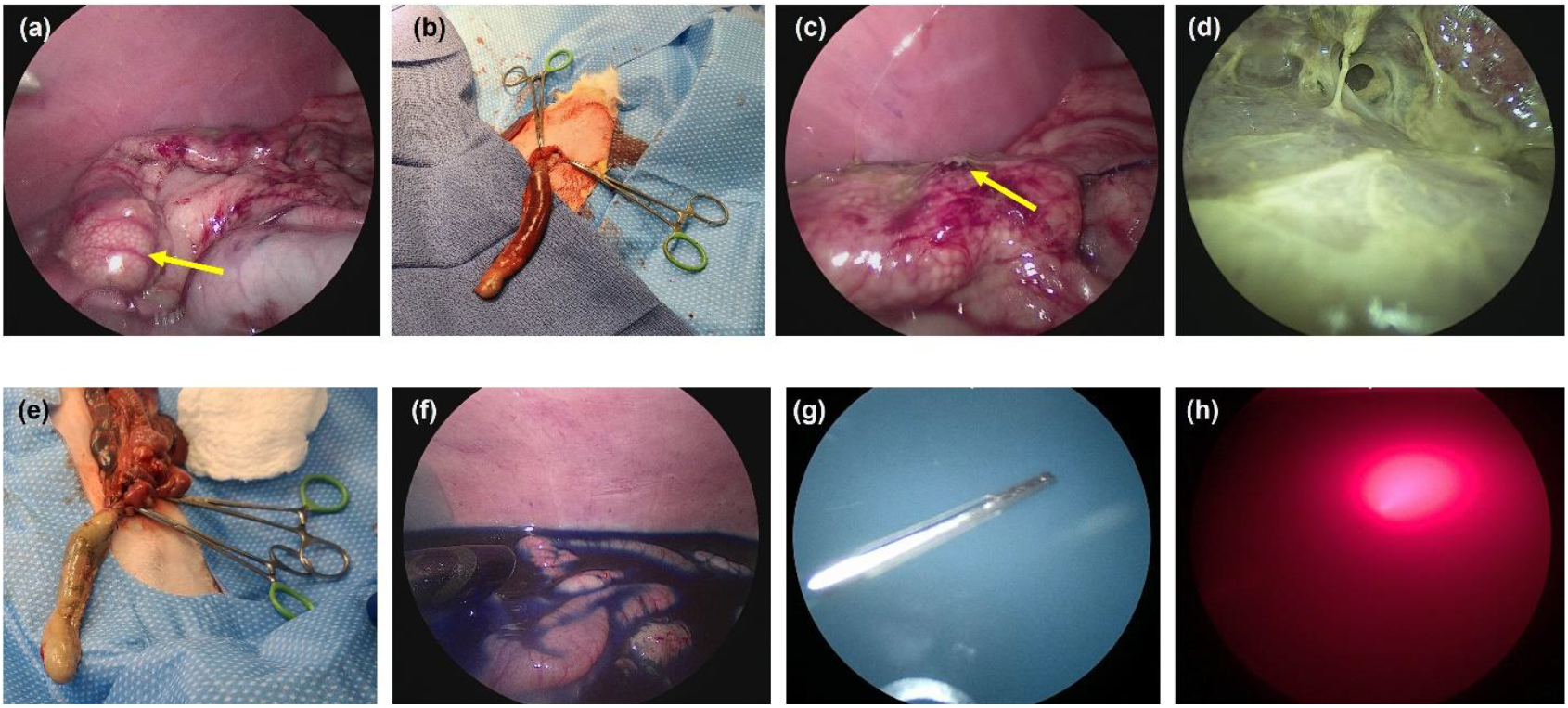
Representative photos of the rabbit model of perforated appendicitis. (a) Appendix pre-ligation (yellow arrow); (b) Ligated appendix immediately following ligation; (c) Appendiceal perforation (yellow arrow) 24 hours post-ligation; (d) feculent peritonitis 24 hours post-ligation; (e) Necrotic appendix 24 hours post-ligation; (f) Methylene blue incubation; (g) Introduction of laser fiber into diluted Intralipid; (h) Laser light delivery.

### In vivo photodynamic therapy and control condition

For full details of the experimental conditions, please see the Supplemental Material. The time from Day 1 to Day 2 procedures ranged from 24 to 40 hours during model development. On Day 2, diagnostic laparoscopy was performed to confirm the presence of perforation and feculent peritonitis (Figure 2d). Peritoneal aspirates were obtained using sterile laparoscopic suction. Then, via laparotomy, the appendix was identified (Figure 2e) and an appendectomy was performed by placing a second monofilament suture ligature at the appendiceal base and dividing the appendix between the sutures using electrocautery. The laparotomy incision was then re-closed in layers, leaving a 5-mm defect to accommodate a 5-mm laparoscopic trocar. Rabbits then either received PDT (n=9) or control conditions (n=10).

For PDT animals, the peritoneal cavity was filled warm, sterile MB (Figure 2f), which was allowed to incubate for 10 minutes to enable bacterial uptake while minimizing uptake by host tissues^32^. Following MB aspiration and flushing of the peritoneal cavity with saline, the abdomen was filled with a sterile fat emulsion to serve as a light-scattering vehicle (Figure 2g). An optical fiber was inserted under laparoscopic visualization to the approximate center of the abdominal cavity (Figure 2g) with care taken to avoid direct contact of the laser fiber with the bowel. Laser light (665 nm, irradiance 20 mW/cm^2^) was delivered to a desired fluence of 25 J/cm^2^ (Figure 2h), resulting in approximately 20 minutes of illumination. For the first 5 PDT rabbits, the laser fiber was manually held – taking care to maintain the fiber stationary in the center of the cavity during the 20-minute light delivery period. In the remaining PDT-treated rabbits, a prototype laparoscopic device was used to introduce and stabilize the optical fiber during treatment (Figure 3)^33^. After illumination, aspiration of the fat emulsion and flushing of the peritoneal cavity was performed. Control animals, representing a drug-only (MB) condition, received all procedures described above, but did not receive laser illumination. Animals were then returned to the vivarium for 24 hours.

**Figure 3:**
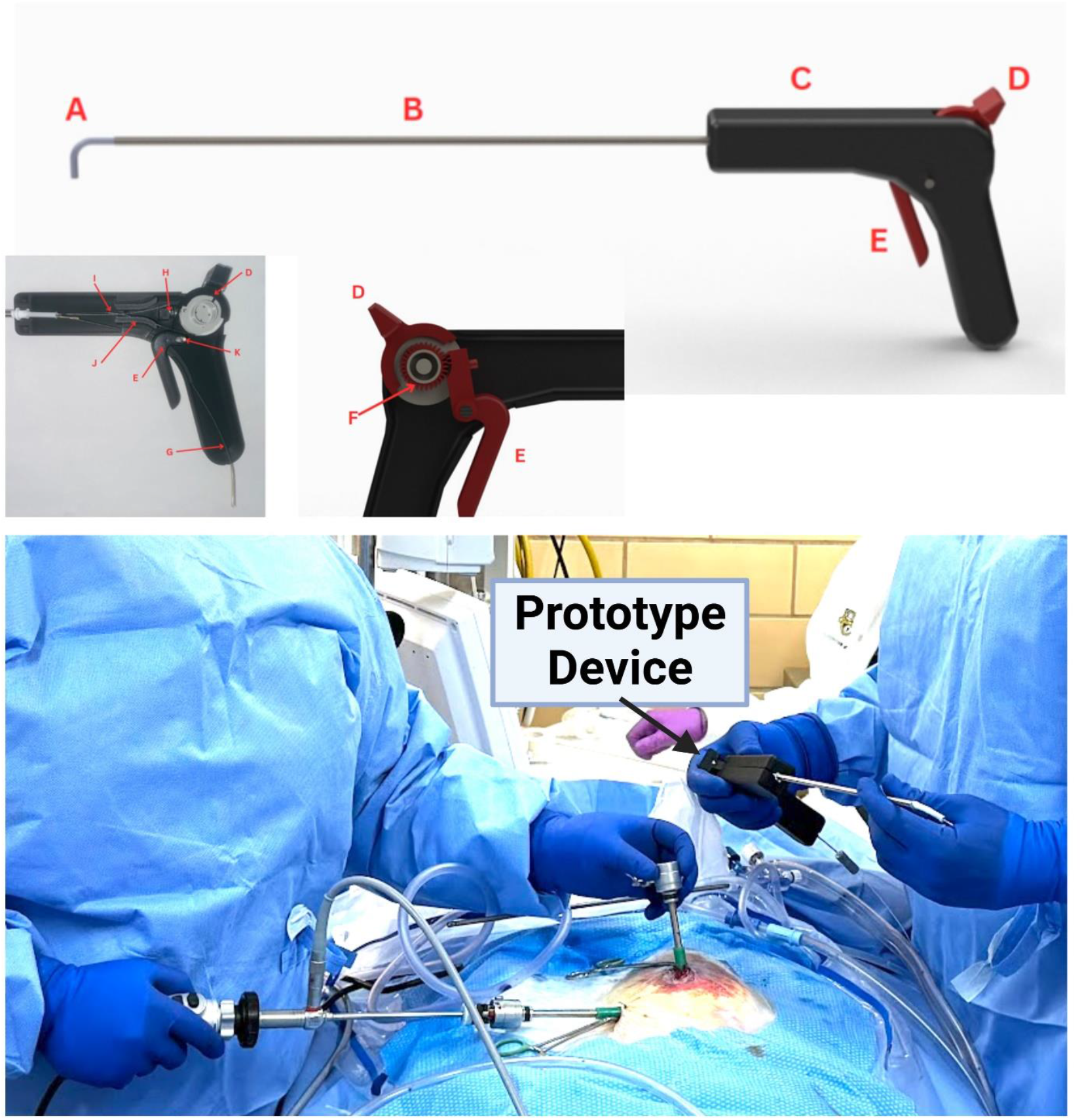
Laparoscopic device for PDT delivery. (top) Rendering of prototype laparoscopic device. Primary components include **(A)** an articulating tip, **(B)** a stiff laparoscopic shaft, and **(C)** a handle. Central tubing **(G)** allows fiberoptic delivery. The user grasps and operates the handle with a single hand, while controlling tip articulation with a wheel knob **(D)** and ratcheting trigger components **(E and F)**. (bottom) Intraoperative deployment of the prototype laparoscopic device during a representative rabbit PDT procedure.

### Collection of post-intervention samples and tissue harvesting

Animals were sacrificed on Day 3, 24 hours after PDT or control conditions. Immediately following sacrifice, a post-intervention peritoneal aspirate was collected (sterile collection under laparoscopic guidance). Tissue samples were then collected from the following organs to evaluate off-target effects of intra-abdominal PDT: (1) small intestine, (2) large intestine, (3) bladder, (4) liver, (5) stomach, (6) kidney, and (7) spleen. Tissue samples were placed in 10% formalin for at least 24 hours prior to histopathological evaluation.

### Evaluation of clinical parameters

Rabbit vital signs, including body temperature, heart rate, and respiratory rate, were collected for all animals at five timepoints: (1) pre-ligation on Day 1, (2) intra-ligation on Day 1, (3) post-ligation from Days 1-2, (4) intra-intervention (PDT or control conditions) on Day 2, and (5) post-intervention from Days 2-3. Body weight was also recorded on Days 1 and 2.

### Histopathology for off-target effects

Tissue samples were fixed in 10% neutral buffered formalin for 72 h at 4°C, processed, and embedded in paraffin for sectioning. Five-micron sections were cut and deparaffinized for Hematoxylin and Eosin (H&E) staining. Slides were scanned with the VS110 Virtual Slide Microscope (Olympus, Waltham, MA, USA) for visualization and assessment of off-target effects. A board-certified pathologist, blinded to experimental conditions, then selected two representative sections from each organ for each animal. A total of 64 slides were evaluated. Each slide was assessed for the presence and distribution of inflammation, necrosis, ischemic injury, vascular changes, edema, architectural alterations, microorganisms, and foreign material.

### Bacteriology

Intra-abdominal fluid aspirates were plated on BHI (Criterion Brain Heart Infusion (BHI)) Broth with 1.5% BD Bacto Agar, Difco MacConkey agar, and blood agar (Difco Columbia Broth with 1.5% BD Bacto Agar and 5% HemoStat Laboratories Defib Sheep Blood). Gram stain, oxidase, and indole tests were performed to initially classify the bacteria. Cultures were incubated aerobically (except for the blood agar cultures which were incubated in candle jars) at 37ºC for 24 hours and bacterial growth was quantified in units of colony-forming units (CFU) per mL. Genomic DNA samples were extracted from bacterial isolates as described in the Supplemental Material.

### In vitro photodynamic therapy

For *in vitro* experiments, bacteria were grown overnight on an orbital shaker at 37ºC in 25 mL of BHI. Bacteria were harvested the next day by centrifugation at 5000xg for 15 minutes at 4 ºC and washed twice with 25 mL of PBS. Samples were resuspended at an optical density of 1 at 600 nm. Eight 1 mL aliquots were made using diluted samples. Four aliquots were incubated with MB (MB+) (Akorn, Inc., Lake Forest, IL) at a concentration of 300 µg/mL for 30 minutes at room temperature in a dark room on a rotating mount, and the remaining four with an equivalent volume of PBS (MB-). Following incubation, the samples were centrifuged at 10,000xg for 10 minutes, washed twice with PBS using the same centrifugation settings, and then transferred to 12-well tissue culture dishes for PDT experiments. Illumination was delivered by a lens-tipped optical fiber coupled to a 665 nm laser (LDX-3230-665, LDX Optronics, Inc., Maryville, TN) at an irradiance of 4 mW/cm^2^ and fluence of 7.2 J/cm^2^, to replicate previously published conditions^19–21^. Each experiment also included control conditions: drug-only (MB+L-), light-only (MB-L+), and drug-and light-free (MB-L-). Drug-free controls (MB-) were prepared by replacing MB with PBS, as described above. Light-free controls (L-) were placed in the same room as illuminated samples (L+) but shielded from laser light. After PDT or control conditions, suspensions were serially diluted in 96-well tissue culture plates and incubated for approximately 18 hours at 37ºC.

### Statistical Analysis

For *in vivo* experiments, changes in bacterial burden from pre- to post-intervention were compared between PDT and control conditions using the Mann-Whitney test. Longitudinal changes in clinical parameters (weight, temperature, heart rate, respiratory rate) were analyzed using mixed-effects models with the Geisser-Greenhouse correction. Comparisons of clinical parameters between PDT and control conditions at individual timepoints and changes from pre- to post-intervention were performed with the Mann-Whitney test. For *in vitro* experiments, bacterial burden was compared between PDT and control conditions using two-way ANOVA. Reduction in bacterial burden for PDT vs. drug- and light-free controls was compared between Gram-positive and Gram-negative species using the Mann-Whitney test. All statistical analyses were performed using GraphPad Prism (v10.2.1, GraphPad Software, LLC, Boston, MA) and MATLAB (R2022b, The Mathworks, Inc., Natick, MA).

## RESULTS

### Rabbit model of perforated appendicitis

Fourteen of 19 rabbits had PA with feculent peritonitis on Day 2, frank pus was apparent in 3 rabbits, and development of thin fibrin adhesions was observed in 4 rabbits. Three rabbits had no apparent peritonitis or purulent fluid, and 4 died prior to surgery on Day 2. Day 2 peritoneal aspirates were acquired for 2 of 4 rabbits who died before surgery, resulting in a total of 17 pre-intervention aspirate samples. Among the rabbits that survived until surgical intervention on Day 2, clinical signs were consistent with evolving intra-abdominal infection. These animals demonstrated adequate analgesia and resumed oral intake, though with decreased fluid consumption in some cases. Mild-to-moderate abdominal distension was noted, reflecting the presence of peritonitis and/or intra-abdominal inflammation secondary to appendiceal perforation.

Appendiceal necrosis was apparent in all animals, except the last PDT-treated animal, in which the appendiceal artery was not ligated (Figure 2e). However, in this animal, the development of peritonitis was still evident, with abundant murky fluid and fibrinous adhesions observed.

Bacteria isolated from rabbits on Day 2, prior to PDT or control conditions, are tabulated in Supplemental Table 1. The most common isolated species were *Escherichia coli* (58.8%, 10/17), *Enterobacter hormaechei* (29.4%, 5/17), and *Enterococcus faecalis* (*E. faecalis*, 23.5%, 4/17), with only a single sample (5.9%) containing yeast. 64.7% (11/17) of samples contained multiple bacterial species, 35.3% (6/17) had a monomicrobial infection, and 11.8% (2/17) showed no microbial growth.

Of the 19 rabbits, 8 died prior to the study’s natural endpoint. As mentioned above, 4 died between Days 1 and 2 – one due to an apparent mesenteric volvulus with hemorrhagic ascites and diffusely ischemic bowel unrelated to bacterial infection. An additional 3 rabbits died during the Day 2 procedure: 1 death likely due to intraabdominal sepsis, 1 death thought to be more multifactorial, including possible intraabdominal sepsis and prolonged hypoxia related to anesthesia complications, and 1 death due to an evident idiopathic cardiac condition (confirmed on necropsy). One rabbit died between Days 2 and 3, likely due to overwhelming intraabdominal sepsis as the necropsy report did not reveal another etiology of death. Including these rabbit deaths, 7 rabbits received PDT with complete follow-up, and 3 animals received control conditions with complete follow-up. Subsequent results, therefore, focus on these 10 animals. The clinical response to *in vivo* PDT, in terms of temperature, heart rate, and respiratory rate, are shown in Supplemental Material.

### Bacterial response to in vivo photodynamic therapy

Ten of the 19 rabbits had pre- and post-intervention aspirates collected on Days 2 and 3, respectively, 7 received PDT, and 3 received control conditions. The bacterial burden tended to increase for animals receiving PDT compared to control conditions, although this difference was not statistically significant (p = 0.18; Figure 4a). Bacterial burdens for individual animals are shown in Figure 4b. Bacterial burden decreased post-intervention for one PDT-treated animal and increased for all other rabbits examined.

**Figure 4:**
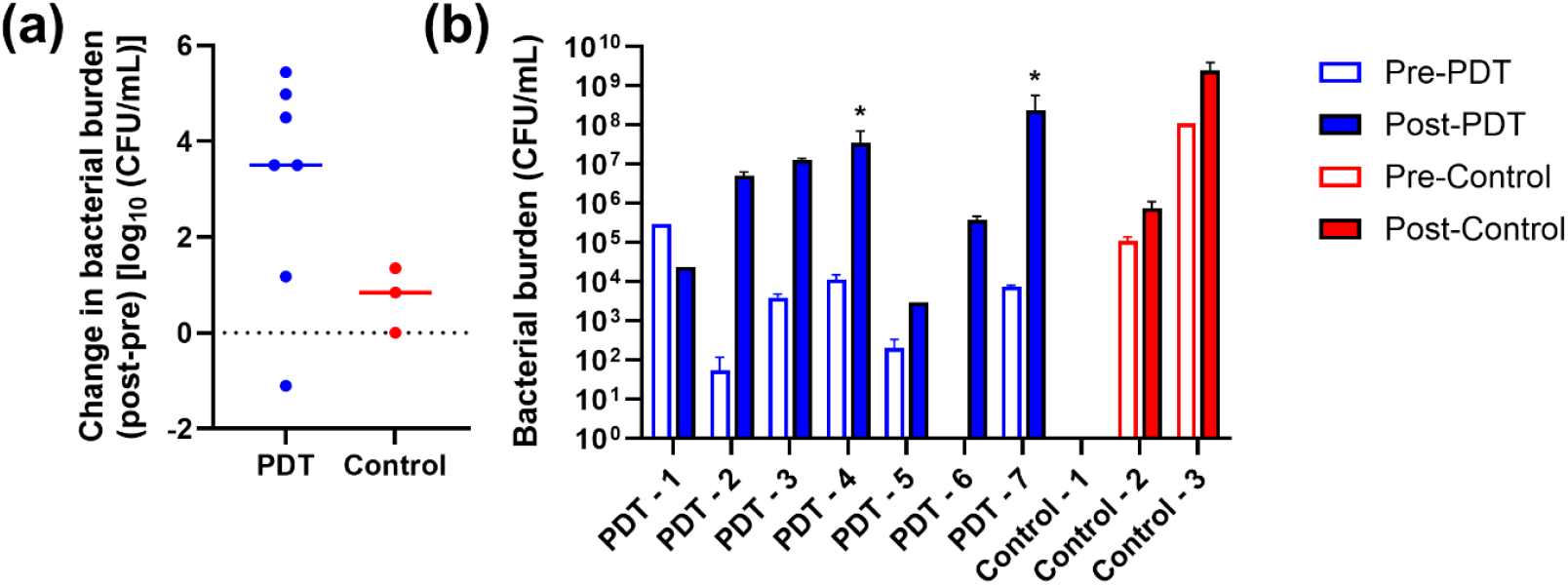
Preliminary *in vivo* PDT efficacy. (a) Change in overall bacterial burden from pre-to post-intervention, for rabbits receiving PDT (n = 7) and control conditions (n = 3). Symbols represent individual animals, with solid horizontal lines representing median values across animals. Horizontal dotted lines indicate no change. (b) Bacterial burden pre- and post-intervention for all animals. Solid bars represent means across media used to culture aspirate, with error bars representing standard deviation. PDT-treated animals in which presumed iatrogenic full-thickness thermal injuries to the small intestine were observed on Day 3 are marked with an asterisk (^*^).

### Evaluation of off-target effects of in vivo PDT

Histologic review of representative slides from intraabdominal organs demonstrated changes consistent with feculent peritonitis in 9 animals, correlating with induced appendiceal perforation in the experimental model. Small intestine perforation, likely due to heat injury, was identified in 1 animal histologically and confirmed on necropsy in 2 animals on Day 3. There were no other histological features specific for MB-PDT-induced injury to the stomach, liver, kidney, small bowel, colon, bladder, or spleen.

In 2 PDT-treated rabbits, at the time of the Day 3 procedure, we observed full-thickness thermal injuries to the small intestine (Supplemental Figure 4), likely due to contact between the optical fiber used for light delivery and the intestine. As these laser-induced perforations likely resulted in bacterial infiltration into the peritoneal cavity unrelated to PA, we re-analyzed the data shown in Figure 4 with these animals removed. Despite removing these animals, the bacterial burden increased for animals receiving PDT relative to control conditions (p=0.39). However, the magnitude of this increase was reduced (2.80±2.74 log_10_ CFU/mL vs. 3.14±2.33 log_10_ CFU/mL).

### In vitro efficacy of PDT against bacteria isolated from a rabbit model of perforated appendicitis

Summary reductions in bacterial burden relative to drug- and light-free controls are shown for Gram-positive and Gram-negative bacteria in Figure 5a, and for individual species in Figure 5b. While PDT significantly reduced the bacterial burden for all species tested (p<0.001), Gram-negative bacteria showed a significantly lower magnitude reduction relative to Gram-positive *E. faecalis* (p<0.001).

**Figure 5:**
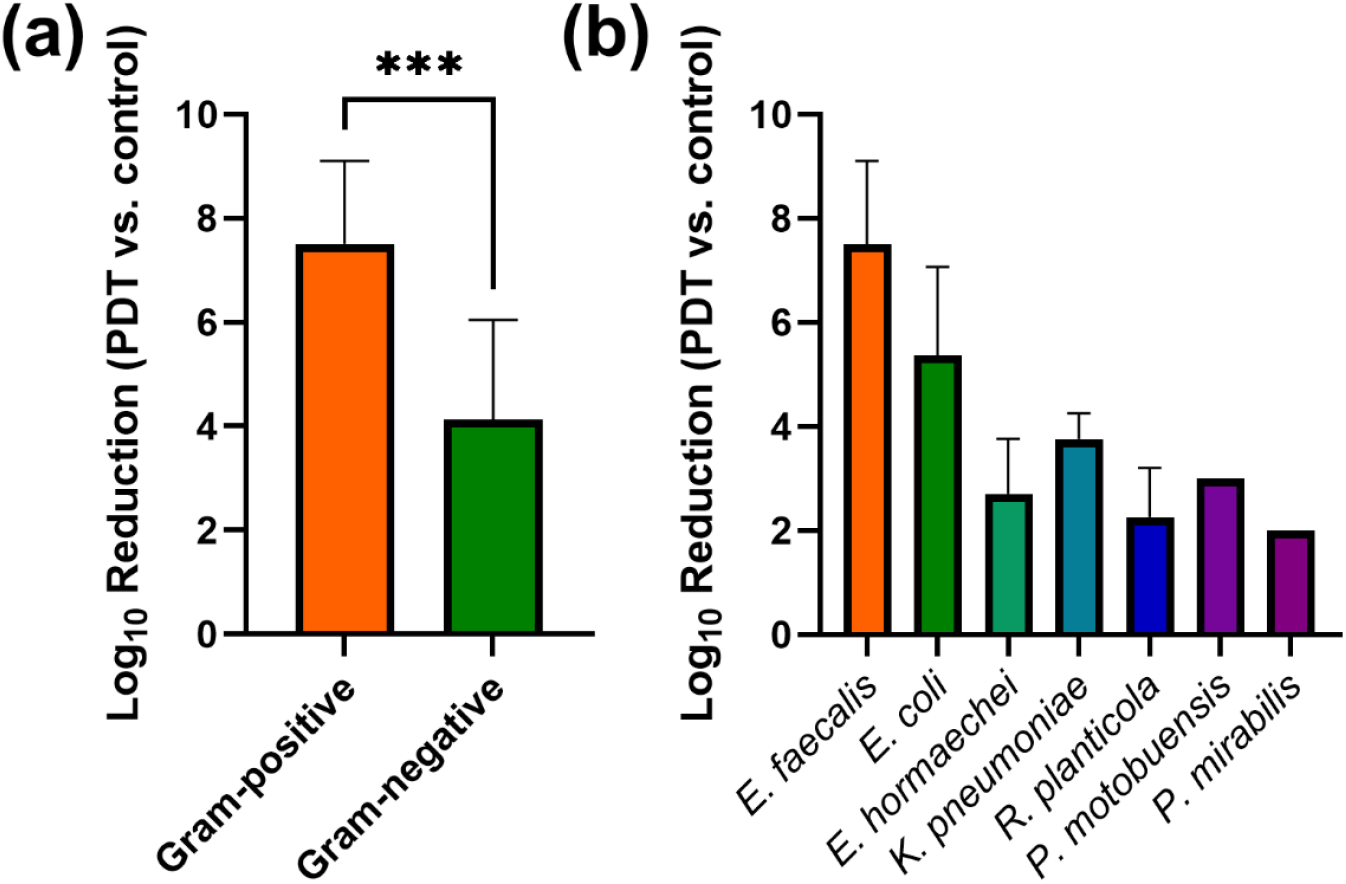
*In vitro* PDT efficacy. Reduction in bacterial burden for PDT-treated samples (MB+L+) relative to drug- and light-free controls (MB-L-), separated by (a) Gram staining and (b) bacterial species. Solid bars indicate mean values across samples, with error bars representing standard deviation across samples. Bacterial species isolated in only a single sample do not include error bars. ^***^ p<0.001

## DISCUSSION

We demonstrated that developing a rabbit model of perforated appendicitis with peritonitis is challenging but feasible. The severity of infection varied substantially across animals despite consistent surgical technique. Laparoscopic delivery of MB-PDT was technically feasible and appeared safe, with no histologic evidence of off-target tissue injury. *In vitro*, MB-PDT achieved >99.9% (3 log_10_) reduction in bacterial burden across all isolated species using a low light dose (4 mW/cm^2^, 7.2 J/cm^2^), indicating that it may be possible to reduce the delivered laser intensity and duration for future *in vivo* studies. Although *in vivo* bacterial reduction was not observed, MB-PDT-treated animals exhibited a trend toward reduced fever, suggesting a partial treatment effect. These results suggest that PDT represents an exciting and potentially valuable alternative adjunctive treatment for intra-abdominal infection secondary to PA and other sources of intra-abdominal contamination. Additional investigation, including optimal light delivery, dosage, and device development are required prior to translation in humans.

The discrepancy between *in vitro* and *in vivo* efficacy likely reflects challenges inherent to light delivery *in vivo*. Delivery of a uniform light dose to the site of infection within the peritoneal cavity is difficult due to anatomic variability and limitations in fiber positioning. The delivered irradiance at the abdominal wall varies nonlinearly with distance from the light source, and absorption depends on tissue optical properties^34,35^. An additional consideration is that the interval between induction of infection and treatment (24–40 hours) may not have allowed for sufficient bacterial proliferation in some animals. Notably, the one animal with the highest pre-treatment burden (figure 4B) was the only case with post-PDT reduction.

Multiple groups have reported rabbit models of appendicitis using appendiceal ligation^28–31^, but we observed wider variability in infection severity, despite consistent surgical technique, animal sources, and housing conditions. Infection severity led to early mortality in 6 of 19 rabbits; 2 animals died for reasons seemingly unrelated to infection (mesenteric volvulus, idiopathic cardiac condition). Additionally, unlike prior reports, partial thickness cautery burn of the distal appendix tip did not result in a reproducible perforation over 24 hours. In two animals, partial thickness burns healed without perforation, prompting a transition to full-thickness cautery perforations. We also found that omitting appendiceal artery ligation resulted in peritonitis without appendiceal necrosis, potentially offering a more clinically relevant model, while avoiding the presence of a dead organ in the peritoneal cavity and the associated intraabdominal sepsis burden.

This study has several limitations. Given the variability and high mortality observed in the rabbit model, alternative small animal models of intra-abdominal infection may offer advantages. Rodent models using cecal ligation and puncture, bacterial inoculation, or fecal slurry have demonstrated reproducible infection and allow for higher-throughput study, though they do not fully replicate the inflammatory profile of PA^36,37^. Mortality prior to intervention reduced the final sample size and limited statistical power, making it difficult to provide conclusive evidence of the *in vivo* PDT anti-bacterial effect. Additionally, only a single combination of drug and light parameters was examined *in vivo*. As we observed high *in vitro* efficacy at lower light doses, it is possible that the dose combination was not optimal. However, similar drug and light parameters showed good efficacy in a recent Phase I clinical trial of intraabdominal abscesses in humans^17^. Future work in small animal models with larger sample sizes will allow more rigorous evaluation of MB-PDT dosing and delivery strategies.

Despite these limitations, this study establishes the feasibility of laparoscopic MB-PDT for intra-abdominal infection and confirms strong *in vitro* efficacy against clinically relevant pathogens. These findings warrant continued investigation of MB-PDT as a novel intraoperative antimicrobial strategy.

## ACKNOWLEDGEMENTS

This work was funded by grant AI178152 from the National Institutes of Health.

