## Supplemental Material for "Intra-Abdominal Photodynamic Therapy in a Rabbit Model of Perforated Appendicitis"

### METHODS

*Induction of perforated appendicitis*

On Day 1, rabbits were anesthetized with ketamine (15-20 mg/kg) plus dexmedetomidine (0.05-0.1 mg/kg), maintained on isoflurane inhalation (1-2%), and hemodynamically monitored throughout the surgery. After shaving the abdomen, the abdomen was prepped and draped in the usual fashion. An approximate 1.5 cm vertical incision was made in the epigastrium. This was carried through the muscle layers and peritoneum sharply. Once identified, the colon was delivered through the ventral incision. The rabbit cecum was identified and followed to the appendix (Figure 2a,b). The perforator vessels within the meso-appendix were cauterized and divided distal-to-proximal. At approximately the appendiceal base (~10 cm from the tip), the appendiceal artery was identified tactilely and ligated via suture ligation. Appendiceal artery ligation was intentionally not performed in the last treated animal. Once hemostasis was confirmed, the appendix was ligated at its base using a 1-0 absorbable monofilament (polydioxanone [PDS]) pre-looped ligature (ENDOLOOP Ligature, Ethicon US, LLC, Raritan, NJ) (Figure 2b). A full-thickness linear burn was then created on the anti-mesenteric surface near the appendiceal base using electrocautery. The appendiceal perforation was confirmed via expression of fecal matter from the burn site. The appendix and cecum were then placed back into the abdomen, taking care to avoid volvulus. The abdominal fascia was closed with braided absorbable suture (polyglactin 910) in a running fashion, and the skin was closed with an interrupted absorbable monofilament suture (poliglecaprone 25) in a horizontal mattress fashion. The incision was then covered with surgical glue (VetBond, 3M), and a weight-based dose of 1% bupivacaine was injected at the incision site as local anesthetic. The rabbit was then allowed to awaken and recover. The rabbit was then returned to the vivarium for 24 hours, during which time perforated appendicitis with diffuse peritonitis developed (Figure 2c). Multimodal analgesia was provided to all rabbits, which included a transdermal fentanyl patch (12-25 µg/hour) and nonsteroidal anti-inflammatory drugs.

*In vivo photodynamic therapy and control condition*

On Day 2, each rabbit was again anesthetized with isoflurane. The time from Day 1 to Day 2 procedures ranged from 24 to 40 hours during model development. Under sterile conditions, a 5-mm laparoscopic trocar was inserted into the superior aspect of the previous laparotomy incision, and the abdomen was insufflated with $CO_2$ to a pressure of 4 mmHg. A 5-mm, 30-degree laparoscope was introduced into the abdomen to inspect and confirm the presence of perforation and feculent peritonitis (Figure 2d). Peritoneal aspirates were obtained using laparoscopic suction, evaluated as described below, and further used for in vitro evaluation of PDT efficacy. Once peritoneal aspirates were obtained, the previous laparotomy incision was reopened. The colon and cecum were delivered via the ventral incision. The appendix was identified (Figure 2e) and an appendectomy was performed using a second PDS pre-looped ligature at the appendiceal base approximately 1 cm from the first. The appendix was divided between the two ligatures using electrocautery. After inspecting the remaining enteric contents for injury and hemostasis, the colon and cecum were delivered back into the abdomen. The midline fascial incision was closed in a running fashion as described on Day 1. However, a 5-mm defect was left to accommodate a single 5-mm laparoscopic trocar. Following sample collection and appendectomy, rabbits either received PDT (n=9) or control conditions (n=10).

For animals receiving PDT, the entire peritoneal cavity was filled with 500 mL of 300 µg/mL (938 µM) warm, sterile methylene blue (MB, Akorn, Inc., Lake Forest, IL), using a laparoscopic suction-irrigator to distribute the liquid throughout the abdomen (Figure 2f). MB was allowed to incubate for 10 minutes to enable bacterial uptake while minimizing uptake by host tissues33. The laparoscopic suction irrigator was then used to aspirate MB, and the peritoneal cavity was flushed twice with 500 mL of warm, sterile saline. The abdomen was irrigated with saline until the effluent returned clear. The remaining saline was aspirated, and the abdomen was then filled with 500 mL of a 0.1% warm, sterile fat emulsion (Intralipid, Baxter Healthcare Corporation, Deerfield, IL) to serve as a vehicle that scatters treatment light throughout the peritoneal cavity (Figure 2g). Immediately following this, a sterile optical fiber (Varilase Platinum Bright Tip, Vascular Solutions, Inc., Minneapolis, MN) was inserted under laparoscopic visualization to the approximate center of the abdominal cavity (Figure 2g). Care was taken to avoid direct contact of the laser fiber with the bowel. Laser light at 665 nm (ML7710, Modulight Corporation, Tampere, Finland) was then delivered at an irradiance of 20 $mW/cm^2$ to a desired fluence of 25 $J/cm^2$ (Figure 2h), resulting in approximately 20 minutes of illumination. For the first five rabbits receiving PDT, the laser fiber was manually held – taking care to maintain the fiber stationary in the center of the cavity during the 20-minute light delivery period. In the remaining PDT-treated rabbits, a prototype laparoscopic device was used to introduce the optical fiber and keep it in in place during treatment (Figure 3). After illumination, the optical fiber was removed from the abdomen, and Intralipid was aspirated. The peritoneal cavity was again flushed twice with warm, sterile saline, until the effluent returned clear. The remaining saline was aspirated. Control animals received all procedures described above, but did not receive laser illumination. These controls, therefore, represent a drug-only (MB) condition.

After delivery of either PDT or control conditions, the laparoscopic trocars were withdrawn from the rabbit's abdomen and the 5 mm fascia at all port sites was closed using absorbable braided suture. The skin was closed using absorbable monofilament suture and covered with surgical glue, as described above. Anesthesia was stopped and the rabbit was extubated at the end of the case. The rabbit was then returned to the vivarium for 24 hours.

*Collection of post-intervention samples and tissue harvesting*

Animals were sacrificed on Day 3, 24 hours after PDT or control conditions. Immediately following sacrifice, a post-intervention peritoneal aspirate was collected using the same procedure as the pre-intervention sample collection on Day 2 (sterile collection under laparoscopic guidance). For animals with scant intra-abdominal fluid present, the peritoneal cavity was rinsed with sterile saline. This saline was then aspirated to serve as the post-intervention sample.

After sterile collection of post-intervention aspirates, the laparoscopic trocar was removed from the abdomen, and a midline laparotomy was performed. Tissue samples were collected from the following organs to evaluate off-target effects of intra-abdominal PDT: (1) small intestine, (2) large intestine, (3) bladder, (4) liver, (5) stomach, (6) kidney, and (7) spleen. Tissue samples were placed in 10% formalin for at least 24 hours before the histopathological evaluation described below.

*Genomic DNA samples*

Genomic DNA samples were extracted from bacterial isolates by suspending a single colony in 0.05 M, pH = 7.5, TRIS buffer in a microcentrifuge tube and adding glass beads (acid-washed, 425-600 μM, Sigma-Aldrich). The tube was inserted into a bead homogenizer (Next Advance Bullet Blender Homogenizer, U.S Patent #5,769,538, Next Advance, NY) for 3 minutes to lyse the bacteria. The mixture was extracted, placed into a clean tube without beads, and stored at -20°C for later use. PCR amplifications of 16S rRNA gene sequences were performed using universal primers: (5' AGA GTT TGA TCC TGG CTC AG3') and (3' ACG GCT ACC TTG TTA CGA CTT) and the iProof PCR kit (Bio-Rad, CA). The thermocycler settings used were the following: denaturation at 98°C for 3 min, 35 cycles [98°C for 60s; 63°C for 45s; 72°C for 45s], and final elongation at 72°C for 5 min. PCR products were purified with the Monarch PCR & DNA Cleanup Kit (New England Biolabs, MA) and sequenced by Plasmidsaurus (Plasmidsauruas, CA) using Oxford Nanopore Technology with custom analysis and annotation. 16S rRNA gene sequences were identified using BLAST[1] with 100% sequence coverage, an E-value of 0.0, with 99-100% identity.

### RESULTS

**Supplemental Table 1:** Microbial species identified from aspirates obtained prior to intervention on Day 2 (n = 17). Values indicate the number of samples that contained each identified species.

| **Bacterial species, No. (%)** | |
|---|---|
| *Escherichia coli* | 10 (58.8%) |
| *Enterobacter hormaechei* | 5 (29.4%) |
| *Enterococcus faecalis* | 4 (23.5%) |
| *Klebsiella pneumoniae* | 3 (17.6%) |
| *Raoultella planticola* | 2 (11.8%) |
| *Bacillus* species | 1 (5.9%) |
| *Paenibacillus motobuensis* | 1 (5.9%) |
| *Proteus mirabilis* | 1 (5.9%) |
| *Candida* species | 1 (5.9%) |

*Clinical response to in vivo photodynamic therapy*

Vital signs (temperature, heart rate, and respiratory rate) were monitored before, during, and after appendiceal ligation, as well as between the ligation procedure and intervention, during the intervention, and post-intervention. Representative time courses for these measures are shown in Supplemental Figure 1, with time measured relative to the beginning of the procedure on each day. Body temperature increased during the post-ligation period, indicating the development of intra-abdominal infection. There was a reduction in temperature during intervention, followed by a post-procedural return to pre-procedure temperature. Over the post-intervention period, temperature showed a larger increase for the control cases, compared to those treated with PDT.

**Supplemental Figure 1: Clinical Parameters.** Longitudinal measurements of (a) temperature, (b) heart rate, and (c) respiratory rate for representative PDT-treated (blue) and control animals (red). Times for pre-ligation and intra-ligation periods are presented relative to the start of the ligation procedure. Times for post-ligation and intra-intervention periods are presented relative to the start of the intervention (PDT or control), and post-intervention times are presented relative to the end of the intervention procedure.

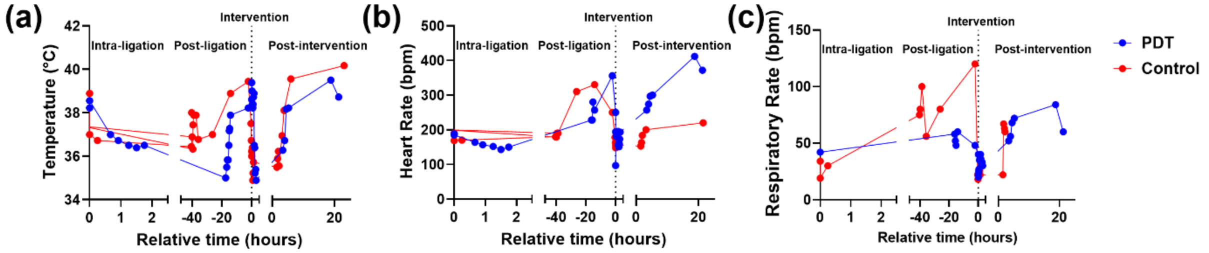


Aggregate results across all animals are shown in Supplemental Figure 2. Longitudinal changes in temperature, heart rate, and respiratory rate were not statistically significant for the PDT or control groups. There were no significant differences between PDT and control at any time point examined. To represent changes in these measurements related to intervention (PDT or control), we examined differences in values from post-ligation to post-intervention (Supplemental Figure 3). There was substantial variability in these quantities, with body temperature displaying the strongest trend from pre- to post-intervention, although this was not statistically significant (PDT: -0.37 ± 1.32 °C, Control: 0.26 ± 0.76 °C; $p = 0.27$).

**Supplemental Figure 2: Summary of Clinical Parameter Results.** Mean values of (a) temperature, (b) heart rate, and (c) respiratory rate across the specified time intervals. Each data point represents the mean value for a particular animal, and horizontal lines represent medians across all animals within a specific group (blue: PDT, red: control).

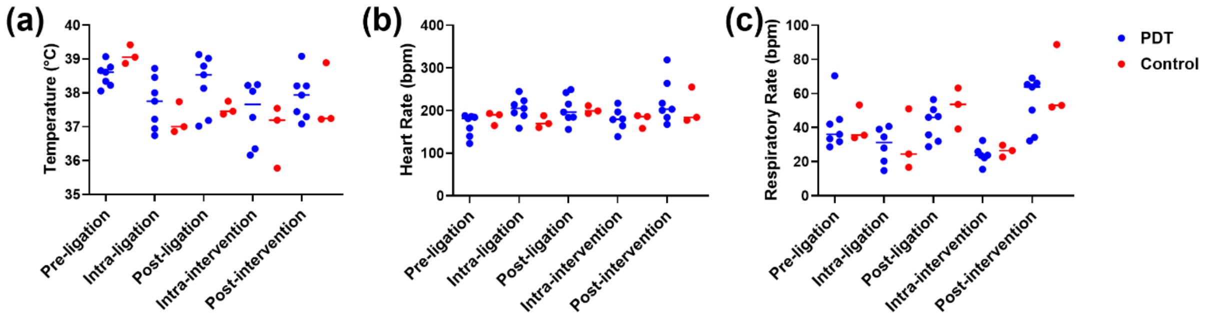

**Supplemental Figure 3: Change in Rabbit Clinical Parameters.** Change in (a) temperature, (b) heart rate, and (c) respiratory rate from the post-ligation to post-intervention period. Individual data points represent mean values for each animal, with horizontal lines representing medians across all animals within a particular group (blue: PDT, red: control). Horizontal dotted lines indicate no change.

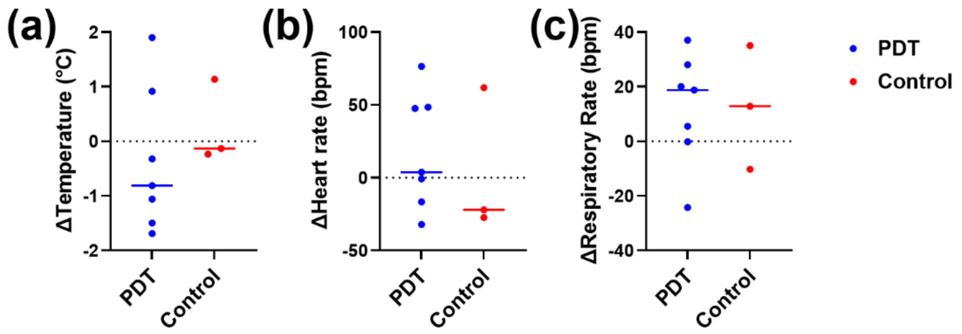


<u>*Evaluation of off-target effects of in vivo photodynamic therapy*</u>

In 2 PDT-treated rabbits, during the Day 3 procedure, we observed full-thickness thermal injuries to the small intestine (Supplemental Figure 4), likely due to contact between the optical fiber used for light delivery and the intestine. As these laser-induced perforations likely resulted in bacterial infiltration into the peritoneal cavity unrelated to perforated appendicitis, we re-analyzed the data shown in Figure 4 with these animals removed. Despite removing these animals, the bacterial burden increased for animals receiving PDT relative to control conditions ($p = 0.39$). However, the magnitude of this increase with reduced (2.80±2.74 $\log_{10}$ CFU/mL vs. 3.14±2.33 $\log_{10}$ CFU/mL).

**Supplemental Figure 4: Bowel perforation induced by direct contact with optical fiber during laser illumination.**

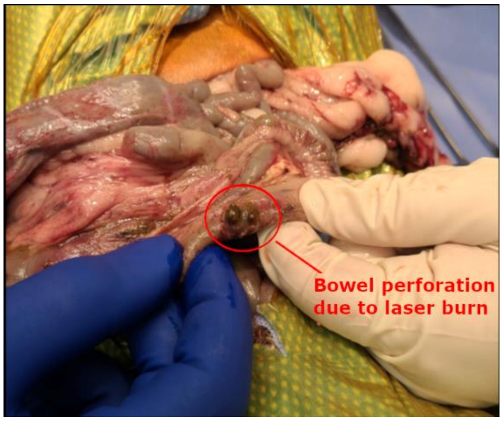


We observed full-thickness thermal injuries to the small intestine in 2 animals, likely due to direct contact between the optical fiber and the small intestine. While the target irradiance was low (20 $mW/cm^2$), direct contact of the optical fiber can result in locally high irradiance, leading to thermal injury. For this reason, we developed and utilized a prototype laparoscopic device to better control the fiberoptic position during laser illumination (Figure 3). This device enabled more reproducible and stable positioning of the fiberoptic tip and reduced hand fatigue for the operator. We anticipate that a similar device could be used for clinical applications after further refinement of the demonstrated prototype.


1 Altschul, S. F., Gish, W., Miller, W., Myers, E. W. & Lipman, D. J. Basic local alignment search tool. Journal of Molecular Biology 215, 403-410, doi:https://doi.org/10.1016/S0022-2836(05)80360-2 (1990).
